# Highly efficient genome editing in barley using novel *Lb*Cas12a variants and impact of sgRNA architecture

**DOI:** 10.1101/2022.04.28.489853

**Authors:** Tom Lawrenson, Alison Hinchliffe, Macarena Forner, Wendy Harwood

## Abstract

We report the first successful, high efficiency use of *Lb*Cas12a in barley and describe the development and application of two novel Cas12a variants. In total we compared five coding sequence (CDS) variants including the two novel ones and two guide architectures over 5 different target genes using twenty different guides. We found large differences in editing efficiencies between the different CDS versions (0-87%) and guide architectures (0-70%) and show our two novel CDS versions massively outperform the others on test in this species. We show heritability of mutations generated. Our findings highlight the importance of optimising CRISPR systems for individual species and are likely to aid the use of *Lb*Cas12a in other monocot species.

## Main text

*Lachnospiraceae bacterium* Cas12a (*Lb*Cas12a) is probably the second most widely used programmable nuclease in plant genome editing after *Streptococcus pyogenes* Cas9 (*Sp*Cas9) and has some potential advantages. Firstly, by nature of its TTTV PAM requirement as opposed to the *Sp*Cas9 requirement of NGG, it has utility in GC deserts which are often found in introns, UTRs and promoter regions. Secondly, *Lb*Cas12a typically produces larger deletions than *Sp*Cas9 which may be useful in deletion studies. Thirdly, whilst *Sp*Cas9 cuts at the PAM proximal end of the target giving blunt ends, *Lb*Cas12a cuts at the PAM distal region, giving sticky ends; two features which may explain the higher incidence of gene targeting achieved with *Lb*Cas12a (Wolter and Puchta, 2019).

Three versions of *Lb*Cas12a, known to function in plants, were tested against one barley target. Firstly, a rice optimised coding sequence (CDS) (*Os*Cas12a) (Tang et al., 2017) secondly a human optimised CDS (*Hs*Cas12a), functional in dicotyledons (Bernabé-Orts et al., 2019), thirdly an *Arabidopsis* optimised CDS containing the D156R “temperature tolerant” mutation (tt*At*Cas12a) (Schindele and Puchta, 2020). We also created two novel versions, *Hs*Cas12a carrying the D156R mutation (tt*Hs*Cas12a) and tt*At*Cas12 carrying 8 introns (tt*At*Cas12+int). These introns had previously boosted *Sp*Cas9 efficiency dramatically (Grutzner 2021) and so we used the same online tool (<u>NetGene2 - 2.42 - Services - DTU Health Tech</u>) to derive a high splicing confidence for the *Arabidopsis* option in our tt*At*Cas12+int design.

The target gene for this comparison was HORVU.MOREX.r3.1HG0069960 using the construct architecture shown in figure 1a. V1 guide architecture was used, after a previous report (Bernabé-Orts et al., 2019) where a single U6 promoter drove a pair of *Lb*Cas12a guides not separated by any additional sequence. *Lb*Cas12a processes a single transcript containing multiple guides into individual guides by recognition of and cleavage at its own DR sequence, which forms the invariable section of guides (Zetsche et al., 2017). A self-processing hepatitis delta ribozyme (HDV) sequence is placed at the 3’ end of the array prior to a terminator to prevent the formation of a spurious additional guide from the final DR. The 5 constructs containing the same V1 4 guide array but five different versions of *LbCas12a* were transformed into barley cultivar Golden Promise using *Agrobacterium* mediated transfer (Hinchliffe and Harwood, 2019) and T0 plants regenerated. DNA was extracted from T0 plants and the HORVU.MOREX.r3.1HG0069960 locus PCR amplified before Sanger sequencing. ABI files were analysed by viewing chromatograms in alignments to wild type sequence using Benchling (https://www.benchling.com/) and targeted mutations were confirmed using the ICE tool (Synthego - CRISPR Performance Analysis) to score plants as either plus or minus for mutagenesis. The number of T0 lines tested/containing mutations is shown in figure 1b. Around 20 T0 lines were created for each of the 5 constructs which showed marked differences in the numbers of lines mutated at the target. *Os*Cas12a showed no mutated lines (0/21). *Hs*Cas12a gave 6/20 (30%) mutated lines which increased to 12/22 (54%) by including the D156 mutation (tt*Hs*Cas12a). The tt*At*Cas12a gave no mutated lines (0/17) but adding introns (tt*At*Cas12a+int) gave 20/23 (87%) mutated lines. This illustrates the huge effect codon usage, the D156 mutation and introns have on the efficiency of *Lb*Cas12a mutagenesis.

**Figure 1.**
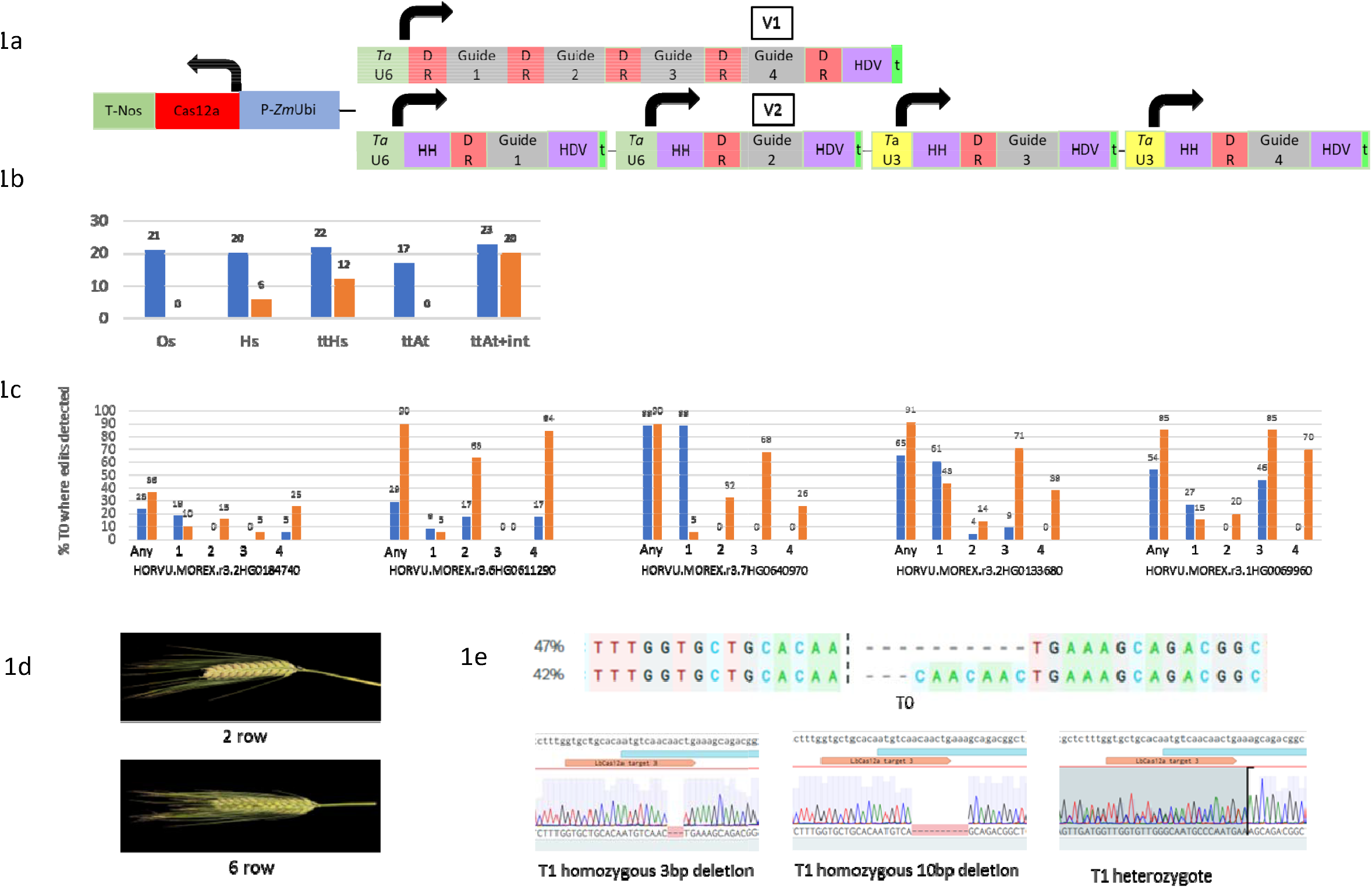
1a. Barley architectures: P-*Zm*Ubi=maize ubiquitin promoter; Cas12a=*Lb*Cas12a CDS; T- Nos=nopaline synthase terminator. *Ta*U6=wheat U6 promoter; *Ta*U3=wheat U3 promoter; DR=direct repeat crRNA; HH/HDV=ribozymes; t=poly-T terminator; V1= V1 array. V2=V2 array. Thick black arrows = direction of transcription. 1b.HORVU.MOREX.r31HG0069960 targeted by V1 4 guide array; different *Lb*Cas12a CDS. *Os*=*Os*Cas12a; *Hs*=HsCas12a; tt*Hs*=tt*Hs*Cas12a; tt*At*=tt*At*Cas12a; tt*At*+int= tt*At*Cas12a+int. Blue bars = number of T0 lines. Orange bars = number of T0 lines containing targeted mutations. 1c. Five barley genes targeted each with tt*Hs*Cas12a/4 guides/V1 vs V2 array. Blue bars = % T0 V1 lines containing targeted mutations. Orange bars = % T0 V2 lines containing targeted mutations. X axis = array guide order. Gene identifiers given. 1d. Left: Wild type 2 row Golden promise. Right: Golden promise T0 plant mutated in HORVU.MOREX.r3.2HG0184740 showing 6 row phenotype. 1e. T0 mutations are inherited in T1 in HORVU.MOREX.r3.1HG0069960.

Although 4 guides were used in the V1 array, only two were active in the *Lb*Cas12a comparison above, so we tried the same 4 guides in a V2 array. This time each guide was driven by a separate TaU6/TaU3 promoter and was flanked by self-cleaving ribozymes; a 5’ Hammerhead (HH) and a 3’ HDV (Wolter 2019). HH/HDV flanked *Lb*Cas12a guides have been shown effective in rice (Zhang et al., 2021). Each HDV was followed by a transcription termination signal to prevent readthrough. This V2 array was coupled with the tt*Hs*Cas12a and used to target HORVU.MOREX.r3.1HG0069960. At the same time 8 further constructs (4 pairs) containing tt*Hs*Cas12a coupled with V1 or V2 guide arrays were made, targeting four further barley genes, each with 4 guides, allowing comparison of V1/V2 guide architectures. Between 19 and 25 T0 lines were created for each construct that were PCR/Sanger sequenced/aligned/ICE tested for targeted mutations. Figure 1c shows the percentage of T0 lines carrying mutations at individual guide targets and the percentage of lines mutated at any guide targets. The V2 array was more efficient overall, giving the greatest percentage of T0 lines mutated at any guide target (36>23; 90>29; 90>88; 91>65; 85>54). This is attributed to guides in array positions 2, 3 and 4 in V2 being more mutagenic than in V1 paired comparisons. It may be that in V1 arrays the single TaU6 promoter can only transcribe short sequences, approximately equivalent in length to a single guide, such that downstream guides in array positions 2,3 and 4 are underrepresented or absent. In V2 arrays, each of the 4 guides may be effectively transcribed due to possessing its own promoter, making guide RNAs in array positions 1-4 abundant. V1 arrays showed higher mutagenesis with guides in array position 1 than V2 comparisons for all five target genes. Guide 1 abundance resulting from the V1 array may be greater than with V2 due to all local transcription factors being available to the single TaU6 promoter in V1. Conversely V2 transcription of guide 1 may be less than in V1 due to local competition for transcription factors between the 4 closely linked TaU6/TaU3 promoters in V2. Reduced promoter activity resulting from the addition of transcription factor binding sites has been reported (Gao 2021).

Knockout of both copies of HORVU.MOREX.r3.2HG0184740 is expected to result in the conversion of two-rowed Golden Promise spikelets into six row spikelets (Komatsuda et al., 2007). This was seen in several active T0 lines as in figure 1d, illustrating the ability of tt*Hs*Cas12a to give knockout phenotypes in the first generation.

The ICE tool calculated one T0 line targeting HORVU.MOREX.r3.1HG0069960 contained 47% & 42% of -10bp & -3bp alleles respectively. From 24 T1 plants, five were T-DNA free, of which two were homozygous for the 3bp deletion, one was homozygous for the 10bp deletion and two were heterozygous (figure 1e). This shows inheritance of mutations detected in T0.

In summary we tested five different *Lb*Cas12a CDS versions and found surprisingly that those codon optimised for rice (*Os*Cas12a) and *Arabidopsis* (tt*At*Cas12a) did not function to a detectable level in barley. A CDS shown to work in various dicot plants, but originally codon optimised for human systems (*Hs*Cas12a) did work in barley and was further improved by the addition of a D156R mutation (tt*Hs*Cas12a). Adding introns to the initially non-functional *Arabidopsis* CDS to give tt*At*Cas12+int transformed it into the most efficient CDS of all in barley. Notably the Netgene splicing tool predictions for *Arabidopsis* proved to be valid in the monocot barley. Our two novel *Lb*Cas12a versions, tt*Hs*Cas12a and tt*At*Cas12a+int both result in highly efficient targeted mutagenesis in barley. We achieved mutagenesis in around 90% of T0 plants for 4/5 barley target genes using tt*Hs*Cas12a/V2 guide architecture but believe this would be increased using tt*At*Cas12a+int which performed best in the Cas12a comparison (87%>54%). We compared two guide architectures and found that V2 gave an overall higher percentage of T0 plants with targeted mutations than V1. Guides in array positions 2,3 and 4 performed hugely better in V2 than in V1. However, guides in array position 1 performed better in V1 than V2.

It is likely that our tt*Hs*Cas12a and tt*At*Cas12a+int CDS modules can increase the reported efficiency of Cas12a mutagenesis in wheat to a similar level seen here in barley. As far as we are aware there is currently only one report of Cas12a use in wheat, where most guides mutated around 5% of their target sites with only one incidence of a substantial increase to 24% (Wang 2021). Our tt*Hs*Cas12a and tt*At*Cas12a+int CDS modules do not require plant growth at the elevated temperatures described in wheat as they both contain the temperature tolerant D156R mutation. The wheat report and another in rice (Zhan 2021) indicate that our V1/V2 guide architecture could be improved upon by using polymerase II promoter driven, single transcript guide arrays, where individual guides are flanked by HH and HDV ribozymes and a polymerase II terminator is used.

## Acknowledgements

We acknowledge support from the project Engineering Nitrogen Symbiosis for Africa (ENSA) currently supported through a grant to the University of Cambridge by the Bill and Melinda Gates Foundation and the Foreign, Commonwealth and Development Office (FCDO). The work was also supported by the UK Biotechnology and Biological Sciences Research Council (BBSRC) through grant BB/P013511/1.

## Author contributions

T.L conceived the ideas, designed the experiments and carried out molecular work. A.H. performed barley transformation. M.F carried out the plant work. T.L. drafted the manuscript. W.H. revised the manuscript.

## Data availability

Component sequences are provided as a supplementary file. Novel coding sequence modules are available through Addgene (www.addgene.org): tt*Hs*Cas12a (Addgene#182385) and tt*AtCas12a*+int (Addgene# 182384). Raw data in the form of ABI files are available from T. Lawrenson.

