## Supplemental information for "Highly efficient genome editing in barley using novel *Lb*Cas12a variants and impact of sgRNA architecture"

**Sequences of construct parts and guides**

TaU6 promoter

crRNA direct repeat

NNNNN: Target specific protospacer sequence

HH ribozyme

HDV ribozyme

Terminator

TaU3 promoter

Nuclear localisation signal

Intron

Maize Ubiquitin promoter

GTGCAGCGTGACCCGGTCGTGCCCCTCTCTAGAGATAATGAGCATTGCATGTCTAAGTTATAAAAAATTACCACATATTTTTTTTGTCACACTTGTTTGAAGTGCAGTTTATCTATCTTTATACATATATTTAAACTTTACTCTACGAATAATATAATCTATAGTACTACAATAATATCAGTGTTTTAGAGAATCATATAAATGAACAGTTAGACATGGTCTAAAGGACAATTGAGTATTTTGACAACAGGACTCTACAGTTTTATCTTTTTAGTGTGCATGTGTTCTCCTTTTTTTTTGCAAATAGCTTCACCTATATAATACTTCATCCATTTTATTAGTACATCCATTTAGGGTTTAGGGTTAATGGTTTTTATAGACTAATTTTTTTAGTACATCTATTTTATTCTATTTTAGCCTCTAAATTAAGAAAACTAAAACTCTATTTTAGTTTTTTTATTTAATAATTTAGATATAAAATAGAATAAAATAAAGTGACTAAAAATTAAACAAATACCCTTTAAGAAATTAAAAAAACTAAGGAAACATTTTTCTTGTTTCGAGTAGATAATGCCAGCCTGTTAAACGCCGTCGACGAGTCTAACGGACACCAACCAGCGAACCAGCAGCGTCGCGTCGGGCCAAGCGAAGCAGACGGCACGGCATCTCTGTCGCTGCCTCTGGACCCCTCTCGAGAGTTCCGCTCCACCGTTGGACTTGCTCCGCTGTCGGCATCCAGAAATTGCGTGGCGGAGCGGCAGACGTGAGCCGGCACGGCAGGCGGCCTCCTCCTCCTCTCACGGCACGGCAGCTACGGGGGATTCCTTTCCCACCGCTCCTTCGCTTTCCCTTCCTCGCCCGCCGTAATAAATAGACACCCCCTCCACACCCTCTTTCCCCAACCTCGTGTTGTTCGGAGCGCACACACACACAACCAGATCTCCCCCAAATCCACCCGTCGGCACCTCCGCTTCAAGGTACGCCGCTCGTCCTCCCCCCCCCCCCCTCTCTACCTTCTCTAGATCGGCGTTCCGGTCCATGGTTAGGGCCCGGTAGTTCTACTTCTGTTCATGTTTGTGTTAGATCCGTGTTTGTGTTAGATCCGTGCTGCTAGCGTTCGTACACGGATGCGACCTGTACGTCAGACACGTTCTGATTGCTAACTTGCCAGTGTTTCTCTTTGGGGAATCCTGGGATGGCTCTAGCCGTTCCGCAGACGGGATCGATTTCATGATTTTTTTTGTTTCGTTGCATAGGGTTTGGTTTGCCCTTTTCCTTTATTTCAATATATGCCGTGCACTTGTTTGTCGGGTCATCTTTTCATGCTTTTTTTTGTCTTGGTTGTGATGATGTGGTCTGGTTGGGCGGTCGTTCTAGATCGGAGTAGAATTCTGTTTCAAACTACCTGGTGGATTTATTAATTTTGGATCTGTATGTGTGTGCCATACATATTCATAGTTACGAATTGAAGATGATGGATGGAAATATCGATCTAGGATAGGTATACATGTTGATGCGGGTTTTACTGATGCATATACAGAGATGCTTTTTGTTCGCTTGGTTGTGATGATGTGGTGTGGTTGGGCGGTCGTTCATTCGTTCTAGATCGGAGTAGAATACTGTTTCAAACTACCTGGTGTATTTATTAATTTTGGAACTGTATGTGTGTGTCATACATCTTCATAGTTACGAGTTTAAGATGGATGGAAATATCGATCTAGGATAGGTATACATGTTGATGTGGGTTTTACTGATGCATATACATGATGGCATATGCAGCATCTATTCATATGCTCTAACCTTGAGTACCTATCTATTATAATAAACAAGTATGTTTTATAATTATTTTGATCTTGATATACTTGGATGATGGCATATGCAGCAGCTATATGTGGATTTTTTTAGCCCTGCCTTCATACGCTATTTATTTGCTTGGTACTGTTTCTTTTGTCGATGCTCACCCTGTTGTTTGGTGTTACTTCTGCAGGA*ATG*

Nopaline synthase terminator

GTCAAGCAGATCGTTCAAACATTTGGCAATAAAGTTTCTTAAGATTGAATCCTGTTGCCGGTCTTGCGATGATTATCATATAATTTCTGTTGAATTACGTTAAGCATGTAATAATTAACATGTAATGCATGACGTTATTTATGAGATGGGTTTTTATGATTAGAGTCCCGCAATTATACATTTAATACGCGATAGAAAACAAAATATAGCGCGCAAACTAGGATAAATTATCGCGCGCGGTGTCATCTATGTTACTAGATCGA

*Os*Cas12a

*ATG*GCTCCTAAGAAGAAGCGGAAGGTTGGTATTCACGGGGTCCCTGCGGCTTCAAAGCTCGAGAAATTCACCAACTGTTATTCGTTGAGCAAAACACTGCGGTTTAAAGCGATTCCAGTCGGCAAGACTCAAGAGAATATAGACAATAAGCGGCTGTTGGTGGAAGATGAAAAGCGCGCGGAGGACTACAAAGGGGTGAAGAAGTTGTTGGACAGATACTACCTCTCTTTTATCAATGATGTCTTGCACTCAATCAAATTGAAGAATCTGAACAACTACATCTCCCTCTTCAGAAAGAAAACAAGGACAGAAAAGGAGAATAAGGAACTTGAAAATTTGGAGATCAATCTGAGGAAAGAGATCGCGAAAGCCTTTAAAGGCAACGAAGGATACAAAAGTCTGTTCAAGAAGGATATAATTGAGACAATTTTGCCAGAGTTCCTCGATGACAAGGACGAGATTGCGCTGGTCAATTCGTTCAACGGATTCACAACAGCATTCACAGGCTTCTTTGATAATCGGGAAAATATGTTCTCTGAGGAGGCAAAGTCCACTTCTATTGCGTTCAGGTGTATCAATGAGAATCTCACTAGGTACATTTCCAACATGGATATCTTTGAGAAGGTTGACGCAATTTTTGACAAGCACGAAGTTCAGGAGATTAAGGAGAAGATCCTCAATTCCGATTATGACGTTGAGGACTTCTTCGAAGGTGAGTTTTTTAATTTCGTGCTCACTCAAGAGGGTATCGACGTGTATAATGCGATCATCGGTGGGTTCGTGACTGAGTCCGGTGAAAAGATTAAGGGATTGAACGAGTATATCAACCTTTACAACCAAAAGACGAAACAGAAGCTGCCAAAGTTCAAGCCTCTTTACAAACAGGTTCTTTCAGACCGCGAGTCACTCTCGTTCTATGGGGAGGGCTACACTTCGGATGAGGAAGTCCTGGAGGTGTTCAGGAATACTCTCAATAAGAATTCGGAGATTTTCTCTTCTATAAAAAAACTGGAAAAGTTGTTTAAGAATTTTGACGAATACTCTAGCGCCGGCATATTTGTGAAAAACGGCCCGGCCATATCAACGATAAGTAAAGATATCTTCGGCGAATGGAACGTGATCAGAGACAAATGGAACGCGGAGTATGACGATATTCACCTGAAGAAGAAGGCTGTCGTAACGGAGAAGTACGAGGATGATCGCAGGAAAAGCTTCAAAAAGATCGGAAGTTTCAGCCTGGAACAGTTGCAGGAGTATGCTGACGCCGATCTTAGCGTCGTCGAGAAGTTGAAGGAGATAATCATCCAAAAGGTCGACGAGATATATAAAGTCTATGGATCAAGTGAAAAACTGTTCGACGCCGACTTCGTTTTGGAGAAGTCCCTGAAGAAGAACGACGCTGTTGTTGCCATTATGAAGGATCTGCTCGACAGCGTGAAGAGTTTCGAGAACTATATTAAGGCTTTTTTCGGGGAGGGGAAGGAGACTAACAGAGATGAGTCCTTCTACGGAGACTTCGTCCTCGCGTACGATATACTCCTTAAGGTAGACCACATCTACGACGCAATCAGAAATTACGTGACACAAAAGCCGTACAGCAAGGACAAGTTCAAACTCTACTTCCAGAACCCCCAGTTCATGGGCGGCTGGGACAAGGACAAGGAAACGGATTACAGGGCTACGATCCTGAGGTATGGTTCAAAATACTACTTGGCGATTATGGACAAGAAGTACGCCAAGTGTCTCCAGAAGATTGACAAAGACGATGTCAATGGCAATTATGAGAAGATCAACTACAAGCTGCTTCCGGGTCCGAACAAGATGCTCCCAAAGGTTTTCTTCAGCAAGAAATGGATGGCCTACTATAACCCAAGCGAGGACATCCAGAAGATTTATAAGAACGGTACGTTCAAGAAGGGCGACATGTTCAATCTTAACGACTGTCACAAGCTGATCGACTTCTTCAAAGACTCAATTAGCCGGTACCCAAAGTGGTCTAACGCCTATGACTTCAACTTTTCGGAAACCGAGAAGTACAAGGATATAGCCGGATTTTATAGAGAGGTGGAAGAGCAGGGCTACAAGGTGTCATTCGAGTCCGCCAGCAAGAAGGAAGTGGACAAGCTCGTGGAAGAGGGTAAGCTCTACATGTTCCAGATTTATAATAAAGACTTTAGCGATAAGAGCCACGGGACACCTAATCTCCACACAATGTATTTCAAGCTGCTCTTCGACGAGAATAACCACGGCCAAATCAGGTTGTCAGGAGGGGCTGAACTCTTCATGCGGCGCGCTAGCCTTAAGAAGGAGGAGCTTGTAGTCCACCCTGCGAATAGTCCAATTGCGAATAAGAACCCGGACAATCCTAAAAAGACTACAACATTGAGCTACGACGTGTACAAGGATAAGAGGTTTTCCGAGGATCAGTACGAGCTCCACATCCCGATTGCGATCAACAAGTGCCCAAAGAATATTTTCAAGATAAACACAGAGGTGCGTGTACTCCTGAAGCATGACGACAATCCTTACGTCATTGGGATTGATCGGGGCGAGAGGAACCTCCTCTATATTGTGGTGGTGGACGGGAAGGGGAACATAGTCGAACAGTACTCCCTTAACGAAATAATTAACAATTTCAACGGCATCCGTATCAAGACCGACTACCATTCGTTGCTGGACAAGAAGGAGAAGGAGAGATTTGAGGCGCGGCAAAATTGGACAAGTATCGAGAACATCAAGGAACTCAAAGCAGGTTATATCTCTCAAGTTGTGCATAAGATATGCGAGCTGGTTGAGAAGTATGACGCAGTGATCGCTCTTGAGGACCTCAACTCGGGCTTTAAGAATTCTAGAGTTAAAGTGGAGAAGCAGGTCTATCAAAAGTTCGAGAAGATGCTTATAGATAAGCTCAACTACATGGTCGATAAGAAATCGAACCCATGTGCCACCGGCGGCGCACTCAAAGGTTACCAAATAACAAACAAATTCGAGTCCTTCAAATCGATGAGTACTCAGAATGGGTTCATATTTTATATACCGGCGTGGCTTACGTCTAAGATCGACCCGTCAACTGGTTTTGTCAACCTGTTGAAAACGAAATACACGTCCATTGCCGATTCGAAAAAGTTCATATCTAGTTTTGATCGTATTATGTACGTCCCAGAGGAAGATCTTTTCGAGTTTGCTCTCGACTACAAAAACTTTTCGCGGACCGATGCGGATTACATTAAAAAATGGAAACTCTATTCGTACGGCAACAGAATCAGGATTTTTCGCAACCCTAAGAAGAATAACGTCTTTGATTGGGAGGAAGTTTGCTTGACTAGCGCGTACAAGGAGCTCTTTAATAAGTATGGCATTAACTACCAACAGGGTGATATCAGAGCACTGCTTTGCGAACAATCTGACAAGGCTTTCTACTCATCCTTCATGGCTTTGATGAGCCTGATGCTCCAGATGAGAAATTCAATTACAGGCAGAACCGACGTGGATTTCTTGATCTCCCCGGTTAAAAATTCTGATGGCATCTTTTACGATAGCAGGAACTATGAAGCGCAAGAGAATGCGATTCTGCCAAAAAATGCAGACGCCAACGGTGCCTATAACATCGCCAGGAAAGTCCTGTGGGCGATCGGCCAGTTCAAAAAGGCCGAGGACGAAAAATTGGACAAGGTCAAAATCGCTATCAGCAACAAAGAGTGGCTGGAGTATGCTCAGACATCCGTAAAGCATAAGCGTCCTGCTGCCACCAAAAAGGCCGGACAGGCTAAGAAAAAGAAGTGA

*Hs*Cas12a

*ATG*AGCAAGCTGGAGAAGTTTACAAACTGCTACTCCCTGTCTAAGACCCTGAGGTTCAAGGCCATCCCTGTGGGCAAGACCCAGGAGAACATCGACAATAAGCGGCTGCTGGTGGAGGACGAGAAGAGAGCCGAGGATTATAAGGGCGTGAAGAAGCTGCTGGATCGCTACTATCTGTCTTTTATCAACGACGTGCTGCACAGCATCAAGCTGAAGAATCTGAACAATTACATCAGCCTGTTCCGGAAGAAAACCAGAACCGAGAAGGAGAATAAGGAGCTGGAGAACCTGGAGATCAATCTGCGGAAGGAGATCGCCAAGGCCTTCAAGGGCAACGAGGGCTACAAGTCCCTGTTTAAGAAGGATATCATCGAGACAATCCTGCCAGAGTTCCTGGACGATAAGGACGAGATCGCCCTGGTGAACAGCTTCAATGGCTTTACCACAGCCTTCACCGGCTTCTTTGATAACAGAGAGAATATGTTTTCCGAGGAGGCCAAGAGCACATCCATCGCCTTCAGGTGTATCAACGAGAATCTGACCCGCTACATCTCTAATATGGACATCTTCGAGAAGGTGGACGCCATCTTTGATAAGCACGAGGTGCAGGAGATCAAGGAGAAGATCCTGAACAGCGACTATGATGTGGAGGATTTCTTTGAGGGCGAGTTCTTTAACTTTGTGCTGACACAGGAGGGCATCGACGTGTATAACGCCATCATCGGCGGCTTCGTGACCGAGAGCGGCGAGAAGATCAAGGGCCTGAACGAGTACATCAACCTGTATAATCAGAAAACCAAGCAGAAGCTGCCTAAGTTTAAGCCACTGTATAAGCAGGTGCTGAGCGATCGGGAGTCTCTGAGCTTCTACGGCGAGGGCTATACATCCGATGAGGAGGTGCTGGAGGTGTTTAGAAACACCCTGAACAAGAACAGCGAGATCTTCAGCTCCATCAAGAAGCTGGAGAAGCTGTTCAAGAATTTTGACGAGTACTCTAGCGCCGGCATCTTTGTGAAGAACGGCCCCGCCATCAGCACAATCTCCAAGGATATCTTCGGCGAGTGGAACGTGATCCGGGACAAGTGGAATGCCGAGTATGACGATATCCACCTGAAGAAGAAGGCCGTGGTGACCGAGAAGTACGAGGACGATCGGAGAAAGTCCTTCAAGAAGATCGGCTCCTTTTCTCTGGAGCAGCTGCAGGAGTACGCCGACGCCGATCTGTCTGTGGTGGAGAAGCTGAAGGAGATCATCATCCAGAAGGTGGATGAGATCTACAAGGTGTATGGCTCCTCTGAGAAGCTGTTCGACGCCGATTTTGTGCTGGAGAAGAGCCTGAAGAAGAACGACGCCGTGGTGGCCATCATGAAGGACCTGCTGGATTCTGTGAAGAGCTTCGAGAATTACATCAAGGCCTTCTTTGGCGAGGGCAAGGAGACAAACAGGGACGAGTCCTTCTATGGCGATTTTGTGCTGGCCTACGACATCCTGCTGAAGGTGGACCACATCTACGATGCCATCCGCAATTATGTGACCCAGAAGCCCTACTCTAAGGATAAGTTCAAGCTGTATTTTCAGAACCCTCAGTTCATGGGCGGCTGGGACAAGGATAAGGAGACAGACTATCGGGCCACCATCCTGAGATACGGCTCCAAGTACTATCTGGCCATCATGGATAAGAAGTACGCCAAGTGCCTGCAGAAGATCGACAAGGACGATGTGAACGGCAATTACGAGAAGATCAACTATAAGCTGCTGCCCGGCCCTAATAAGATGCTGCCAAAGGTGTTCTTTTCTAAGAAGTGGATGGCCTACTATAACCCCAGCGAGGACATCCAGAAGATCTACAAGAATGGCACATTCAAGAAGGGCGATATGTTTAACCTGAATGACTGTCACAAGCTGATCGACTTCTTTAAGGATAGCATCTCCCGGTATCCAAAGTGGTCCAATGCCTACGATTTCAACTTTTCTGAGACAGAGAAGTATAAGGACATCGCCGGCTTTTACAGAGAGGTGGAGGAGCAGGGCTATAAGGTGAGCTTCGAGTCTGCCAGCAAGAAGGAGGTGGATAAGCTGGTGGAGGAGGGCAAGCTGTATATGTTCCAGATCTATAACAAGGACTTTTCCGATAAGTCTCACGGCACACCCAATCTGCACACCATGTACTTCAAGCTGCTGTTTGACGAGAACAATCACGGACAGATCAGGCTGAGCGGAGGAGCAGAGCTGTTCATGAGGCGCGCCTCCCTGAAGAAGGAGGAGCTGGTGGTGCACCCAGCCAACTCCCCTATCGCCAACAAGAATCCAGATAATCCCAAGAAAACCACAACCCTGTCCTACGACGTGTATAAGGATAAGAGGTTTTCTGAGGACCAGTACGAGCTGCACATCCCAATCGCCATCAATAAGTGCCCCAAGAACATCTTCAAGATCAATACAGAGGTGCGCGTGCTGCTGAAGCACGACGATAACCCCTATGTGATCGGCATCGATAGGGGCGAGCGCAATCTGCTGTATATCGTGGTGGTGGACGGCAAGGGCAACATCGTGGAGCAGTATTCCCTGAACGAGATCATCAACAACTTCAACGGCATCAGGATCAAGACAGATTACCACTCTCTGCTGGACAAGAAGGAGAAGGAGAGGTTCGAGGCCCGCCAGAACTGGACCTCCATCGAGAATATCAAGGAGCTGAAGGCCGGCTATATCTCTCAGGTGGTGCACAAGATCTGCGAGCTGGTGGAGAAGTACGATGCCGTGATCGCCCTGGAGGACCTGAACTCTGGCTTTAAGAATAGCCGCGTGAAGGTGGAGAAGCAGGTGTATCAGAAGTTCGAGAAGATGCTGATCGATAAGCTGAACTACATGGTGGACAAGAAGTCTAATCCTTGTGCAACAGGCGGCGCCCTGAAGGGCTATCAGATCACCAATAAGTTCGAGAGCTTTAAGTCCATGTCTACCCAGAACGGCTTCATCTTTTACATCCCTGCCTGGCTGACATCCAAGATCGATCCATCTACCGGCTTTGTGAACCTGCTGAAAACCAAGTATACCAGCATCGCCGATTCCAAGAAGTTCATCAGCTCCTTTGACAGGATCATGTACGTGCCCGAGGAGGATCTGTTCGAGTTTGCCCTGGACTATAAGAACTTCTCTCGCACAGACGCCGATTACATCAAGAAGTGGAAGCTGTACTCCTACGGCAACCGGATCAGAATCTTCCGGAATCCTAAGAAGAACAACGTGTTCGACTGGGAGGAGGTGTGCCTGACCAGCGCCTATAAGGAGCTGTTCAACAAGTACGGCATCAATTATCAGCAGGGCGATATCAGAGCCCTGCTGTGCGAGCAGTCCGACAAGGCCTTCTACTCTAGCTTTATGGCCCTGATGAGCCTGATGCTGCAGATGCGGAACAGCATCACAGGCCGCACCGACGTGGATTTTCTGATCAGCCCTGTGAAGAACTCCGACGGCATCTTCTACGATAGCCGGAACTATGAGGCCCAGGAGAATGCCATCCTGCCAAAGAACGCCGACGCCAATGGCGCCTATAACATCGCCAGAAAGGTGCTGTGGGCCATCGGCCAGTTCAAGAAGGCCGAGGACGAGAAGCTGGATAAGGTGAAGATCGCCATCTCTAACAAGGAGTGGCTGGAGTACGCCCAGACCAGCGTGAAGCACAAAAGGCCGGCGGCCACGAAAAAGGCCGGCCAGGCAAAAAAGAAAAAGTAG

tt*Hs*Cas12a (Addgene#182385)

*ATG*AGCAAGCTGGAGAAGTTTACAAACTGCTACTCCCTGTCTAAGACCCTGAGGTTCAAGGCCATCCCTGTGGGCAAGACCCAGGAGAACATCGACAATAAGCGGCTGCTGGTGGAGGACGAGAAGAGAGCCGAGGATTATAAGGGCGTGAAGAAGCTGCTGGATCGCTACTATCTGTCTTTTATCAACGACGTGCTGCACAGCATCAAGCTGAAGAATCTGAACAATTACATCAGCCTGTTCCGGAAGAAAACCAGAACCGAGAAGGAGAATAAGGAGCTGGAGAACCTGGAGATCAATCTGCGGAAGGAGATCGCCAAGGCCTTCAAGGGCAACGAGGGCTACAAGTCCCTGTTTAAGAAGGATATCATCGAGACAATCCTGCCAGAGTTCCTGGACGATAAGGACGAGATCGCCCTGGTGAACAGCTTCAATGGCTTTACCACAGCCTTCACCGGCTTCTTTCGCAACAGAGAGAATATGTTTTCCGAGGAGGCCAAGAGCACATCCATCGCCTTCAGGTGTATCAACGAGAATCTGACCCGCTACATCTCTAATATGGACATCTTCGAGAAGGTGGACGCCATCTTTGATAAGCACGAGGTGCAGGAGATCAAGGAGAAGATCCTGAACAGCGACTATGATGTGGAGGATTTCTTTGAGGGCGAGTTCTTTAACTTTGTGCTGACACAGGAGGGCATCGACGTGTATAACGCCATCATCGGCGGCTTCGTGACCGAGAGCGGCGAGAAGATCAAGGGCCTGAACGAGTACATCAACCTGTATAATCAGAAAACCAAGCAGAAGCTGCCTAAGTTTAAGCCACTGTATAAGCAGGTGCTGAGCGATCGGGAGTCTCTGAGCTTCTACGGCGAGGGCTATACATCCGATGAGGAGGTGCTGGAGGTGTTTAGAAACACCCTGAACAAGAACAGCGAGATCTTCAGCTCCATCAAGAAGCTGGAGAAGCTGTTCAAGAATTTTGACGAGTACTCTAGCGCCGGCATCTTTGTGAAGAACGGCCCCGCCATCAGCACAATCTCCAAGGATATCTTCGGCGAGTGGAACGTGATCCGGGACAAGTGGAATGCCGAGTATGACGATATCCACCTGAAGAAGAAGGCCGTGGTGACCGAGAAGTACGAGGACGATCGGAGAAAGTCCTTCAAGAAGATCGGCTCCTTTTCTCTGGAGCAGCTGCAGGAGTACGCCGACGCCGATCTGTCTGTGGTGGAGAAGCTGAAGGAGATCATCATCCAGAAGGTGGATGAGATCTACAAGGTGTATGGCTCCTCTGAGAAGCTGTTCGACGCCGATTTTGTGCTGGAGAAGAGCCTGAAGAAGAACGACGCCGTGGTGGCCATCATGAAGGACCTGCTGGATTCTGTGAAGAGCTTCGAGAATTACATCAAGGCCTTCTTTGGCGAGGGCAAGGAGACAAACAGGGACGAGTCCTTCTATGGCGATTTTGTGCTGGCCTACGACATCCTGCTGAAGGTGGACCACATCTACGATGCCATCCGCAATTATGTGACCCAGAAGCCCTACTCTAAGGATAAGTTCAAGCTGTATTTTCAGAACCCTCAGTTCATGGGCGGCTGGGACAAGGATAAGGAGACAGACTATCGGGCCACCATCCTGAGATACGGCTCCAAGTACTATCTGGCCATCATGGATAAGAAGTACGCCAAGTGCCTGCAGAAGATCGACAAGGACGATGTGAACGGCAATTACGAGAAGATCAACTATAAGCTGCTGCCCGGCCCTAATAAGATGCTGCCAAAGGTGTTCTTTTCTAAGAAGTGGATGGCCTACTATAACCCCAGCGAGGACATCCAGAAGATCTACAAGAATGGCACATTCAAGAAGGGCGATATGTTTAACCTGAATGACTGTCACAAGCTGATCGACTTCTTTAAGGATAGCATCTCCCGGTATCCAAAGTGGTCCAATGCCTACGATTTCAACTTTTCTGAGACAGAGAAGTATAAGGACATCGCCGGCTTTTACAGAGAGGTGGAGGAGCAGGGCTATAAGGTGAGCTTCGAGTCTGCCAGCAAGAAGGAGGTGGATAAGCTGGTGGAGGAGGGCAAGCTGTATATGTTCCAGATCTATAACAAGGACTTTTCCGATAAGTCTCACGGCACACCCAATCTGCACACCATGTACTTCAAGCTGCTGTTTGACGAGAACAATCACGGACAGATCAGGCTGAGCGGAGGAGCAGAGCTGTTCATGAGGCGCGCCTCCCTGAAGAAGGAGGAGCTGGTGGTGCACCCAGCCAACTCCCCTATCGCCAACAAGAATCCAGATAATCCCAAGAAAACCACAACCCTGTCCTACGACGTGTATAAGGATAAGAGGTTTTCTGAGGACCAGTACGAGCTGCACATCCCAATCGCCATCAATAAGTGCCCCAAGAACATCTTCAAGATCAATACAGAGGTGCGCGTGCTGCTGAAGCACGACGATAACCCCTATGTGATCGGCATCGATAGGGGCGAGCGCAATCTGCTGTATATCGTGGTGGTGGACGGCAAGGGCAACATCGTGGAGCAGTATTCCCTGAACGAGATCATCAACAACTTCAACGGCATCAGGATCAAGACAGATTACCACTCTCTGCTGGACAAGAAGGAGAAGGAGAGGTTCGAGGCCCGCCAGAACTGGACCTCCATCGAGAATATCAAGGAGCTGAAGGCCGGCTATATCTCTCAGGTGGTGCACAAGATCTGCGAGCTGGTGGAGAAGTACGATGCCGTGATCGCCCTGGAGGACCTGAACTCTGGCTTTAAGAATAGCCGCGTGAAGGTGGAGAAGCAGGTGTATCAGAAGTTCGAGAAGATGCTGATCGATAAGCTGAACTACATGGTGGACAAGAAGTCTAATCCTTGTGCAACAGGCGGCGCCCTGAAGGGCTATCAGATCACCAATAAGTTCGAGAGCTTTAAGTCCATGTCTACCCAGAACGGCTTCATCTTTTACATCCCTGCCTGGCTGACATCCAAGATCGATCCATCTACCGGCTTTGTGAACCTGCTGAAAACCAAGTATACCAGCATCGCCGATTCCAAGAAGTTCATCAGCTCCTTTGACAGGATCATGTACGTGCCCGAGGAGGATCTGTTCGAGTTTGCCCTGGACTATAAGAACTTCTCTCGCACAGACGCCGATTACATCAAGAAGTGGAAGCTGTACTCCTACGGCAACCGGATCAGAATCTTCCGGAATCCTAAGAAGAACAACGTGTTCGACTGGGAGGAGGTGTGCCTGACCAGCGCCTATAAGGAGCTGTTCAACAAGTACGGCATCAATTATCAGCAGGGCGATATCAGAGCCCTGCTGTGCGAGCAGTCCGACAAGGCCTTCTACTCTAGCTTTATGGCCCTGATGAGCCTGATGCTGCAGATGCGGAACAGCATCACAGGCCGCACCGACGTGGATTTTCTGATCAGCCCTGTGAAGAACTCCGACGGCATCTTCTACGATAGCCGGAACTATGAGGCCCAGGAGAATGCCATCCTGCCAAAGAACGCCGACGCCAATGGCGCCTATAACATCGCCAGAAAGGTGCTGTGGGCCATCGGCCAGTTCAAGAAGGCCGAGGACGAGAAGCTGGATAAGGTGAAGATCGCCATCTCTAACAAGGAGTGGCTGGAGTACGCCCAGACCAGCGTGAAGCACAAAAGGCCGGCGGCCACGAAAAAGGCCGGCCAGGCAAAAAAGAAAAAGTAG

tt*At*Cas12a

*ATG*AGCAAGCTCGAGAAGTTTACCAACTGCTACAGCCTCTCTAAGACCCTCAGGTTCAAGGCTATCCCTGTGGGAAAGACCCAAGAGAATATCGACAACAAGAGGCTCCTCGTCGAGGATGAGAAGAGAGCTGAAGATTACAAGGGCGTGAAGAAGCTCCTCGACAGGTACTACCTCAGCTTCATCAACGATGTGCTCCACAGCATCAAGCTCAAGAACCTCAACAACTACATCAGCCTCTTCCGTAAGAAAACCAGGACCGAGAAAGAGAACAAAGAGCTTGAGAACCTCGAGATCAACCTCCGTAAAGAGATCGCCAAGGCTTTCAAGGGAAACGAGGGATACAAGAGCCTCTTCAAGAAGGATATTATCGAGACAATCCTGCCTGAGTTCCTGGACGATAAGGATGAGATCGCTCTCGTGAACAGCTTCAACGGATTCACTACTGCCTTCACCGGATTCTTCAGAAACAGGGAAAACATGTTCAGCGAAGAGGCCAAGAGCACCTCTATCGCTTTCAGATGCATCAACGAGAACCTCACGCGTTACATCAGCAACATGGACATCTTCGAGAAGGTGGACGCCATCTTCGATAAGCACGAGGTGCAAGAAATCAAAGAGAAGATCCTCAACAGCGACTACGACGTCGAGGACTTTTTTGAAGGGGAGTTCTTCAACTTCGTTCTCACCCAAGAGGGCATCGACGTGTACAACGCTATTATCGGAGGATTCGTGACCGAGTCTGGGGAGAAGATTAAGGGACTCAACGAGTACATCAACCTGTACAACCAGAAAACGAAGCAGAAGCTCCCGAAGTTCAAGCCGCTCTACAAGCAGGTTCTCTCTGATCGTGAGAGCCTCTCATTTTACGGTGAGGGTTACACCTCTGACGAGGAAGTGCTTGAGGTTTTCCGTAACACCCTCAACAAGAACAGCGAGATCTTCTCGTCCATCAAGAAGTTGGAGAAGCTTTTCAAGAACTTCGACGAGTACAGCAGCGCTGGGATCTTCGTTAAGAACGGACCTGCTATCAGCACCATCAGCAAGGATATTTTCGGCGAGTGGAACGTGATCAGGGACAAGTGGAATGCTGAGTACGATGACATCCACCTCAAGAAGAAGGCTGTCGTCACTGAGAAGTACGAGGATGACAGGCGTAAGTCGTTCAAGAAGATCGGCTCTTTCAGCCTCGAGCAGCTTCAAGAATACGCTGATGCTGATCTCAGCGTGGTCGAGAAGCTCAAAGAGATCATCATCCAGAAGGTCGACGAGATCTACAAGGTGTACGGGTCCTCTGAGAAGTTGTTCGATGCTGATTTCGTCCTCGAGAAGAGTCTGAAGAAGAACGACGCTGTCGTCGCGATCATGAAGGATTTGCTCGACAGCGTGAAGTCCTTCGAGAACTATATCAAGGCCTTCTTCGGAGAGGGCAAAGAGACTAATAGGGACGAGTCTTTCTACGGGGATTTCGTGCTCGCTTACGATATCCTCCTCAAGGTGGACCATATCTACGACGCCATCAGAAACTACGTGACCCAGAAGCCTTACAGCAAGGACAAGTTCAAGTTGTACTTTCAGAACCCGCAGTTCATGGGCGGATGGGACAAAGACAAAGAGACAGATTACAGGGCCACCATCCTCAGGTACGGGTCTAAGTACTACCTGGCCATCATGGACAAGAAATACGCCAAGTGCCTCCAAAAGATCGACAAGGATGACGTGAACGGGAACTATGAGAAGATCAACTACAAGCTCCTTCCGGGACCGAACAAGATGCTTCCTAAGGTGTTCTTCAGCAAGAAATGGATGGCCTACTACAACCCGTCTGAGGACATCCAGAAAATCTACAAGAACGGGACCTTCAAGAAAGGCGACATGTTCAACCTCAACGACTGCCACAAGCTCATCGATTTCTTCAAGGACAGCATCTCGCGTTACCCGAAGTGGTCTAACGCTTACGACTTTAACTTCAGCGAGACAGAAAAGTACAAGGATATCGCCGGGTTCTACCGTGAGGTTGAGGAACAGGGTTACAAGGTTAGCTTCGAGAGCGCCTCCAAGAAAGAGGTTGACAAGTTGGTCGAAGAGGGCAAGCTCTACATGTTCCAGATCTATAACAAGGACTTCTCCGACAAGAGCCACGGAACTCCTAACCTCCATACGATGTACTTCAAGCTGCTTTTCGACGAGAACAACCACGGGCAGATCAGACTTTCTGGTGGTGCTGAACTCTTCATGCGTAGGGCCTCACTCAAGAAAGAAGAGTTGGTTGTTCACCCGGCCAACTCTCCAATCGCTAACAAGAATCCTGACAACCCGAAAAAGACCACCACGCTGTCTTACGACGTCTACAAGGACAAAAGGTTCAGCGAGGACCAGTACGAGCTTCATATCCCGATCGCTATCAACAAGTGCCCGAAGAACATCTTCAAGATCAATACCGAGGTGAGGGTGCTGCTCAAGCACGATGATAACCCTTACGTGATCGGAATCGATCGTGGTGAGAGAAACCTCCTCTACATCGTTGTGGTGGACGGAAAGGGAAACATCGTCGAGCAGTACAGCCTGAACGAGATTATCAACAATTTCAACGGCATCAGGATCAAGACCGACTACCACTCACTCCTCGATAAGAAAGAAAAAGAGCGTTTCGAGGCCAGGCAGAACTGGACTTCTATCGAAAACATCAAAGAGTTGAAGGCCGGCTACATCTCTCAGGTGGTGCATAAGATCTGCGAGCTGGTGGAAAAGTACGATGCTGTGATCGCTCTTGAGGACCTCAACTCTGGGTTCAAGAACAGTAGAGTGAAGGTTGAGAAGCAGGTCTACCAAAAGTTCGAGAAGATGCTCATCGACAAGCTCAACTACATGGTGGACAAAAAGAGCAACCCTTGCGCTACCGGTGGTGCTCTTAAGGGATACCAGATCACGAACAAGTTCGAGTCCTTCAAGAGCATGAGCACCCAGAACGGCTTCATCTTCTATATCCCTGCTTGGCTCACCAGCAAGATCGATCCTTCTACTGGTTTCGTGAACCTGCTCAAGACCAAGTACACCTCGATCGCCGACAGCAAGAAGTTCATCTCGTCTTTCGACAGGATCATGTACGTGCCGGAAGAGGATCTTTTCGAGTTCGCTCTCGACTATAAGAACTTCAGCAGGACCGACGCCGACTACATTAAGAAGTGGAAGCTCTACTCCTACGGGAACCGTATCAGGATCTTCCGAAATCCGAAGAAAAACAACGTGTTCGACTGGGAAGAAGTGTGCCTCACCTCTGCCTACAAAGAACTGTTCAACAAGTACGGCATCAACTACCAGCAGGGTGATATCAGGGCTCTTTTGTGCGAGCAGAGCGACAAGGCATTCTACAGCTCATTCATGGCCCTCATGTCTCTCATGCTCCAGATGAGGAACTCTATCACCGGAAGGACCGATGTGGACTTCCTTATCTCTCCGGTCAAGAACTCTGACGGGATCTTCTACGACAGCCGTAACTATGAGGCTCAAGAGAACGCTATCCTGCCGAAGAATGCTGATGCAAACGGGGCTTACAACATTGCGAGAAAGGTTCTCTGGGCTATCGGGCAGTTTAAGAAAGCGGAAGATGAGAAGCTGGACAAGGTGAAGATCGCCATCTCCAACAAAGAGTGGCTTGAGTACGCTCAGACCTCCGTTAA

tt*At*Cas12a+int (Addgene# 182384)

*ATG*AGCAAGCTCGAGAAGTTTACCAACTGCTACAGCCTCTCTAAGACCCTCAGGTTCAAGGCTATCCCTGTGGGAAAGACCCAAGAGAATATCGACAACAAGAGGCTCCTCGTCGAGGATGAGAAGAGAGCTGAAGATTACAAGGGCGTGAAGAAGCTCCTCGACAGGTACTACCTCAGCTTCATCAACGATGTGCTCCACAGCATCAAGCTCAAGAACCTCAACAACTACATCAGCCTCTTCCGTAAGAAAACCAGGACCGAGAAAGAGAACAAAGAGCTTGAGAACCTCGAGATCAACCTCCGTAAAGAGATCGCCAAGGCTTTCAAGGGAAACGAGGGATACAAGAGCCTCTTCAAGAAGGATATTATCGAGACAATCCTGCCTGAGTTCCTGGACGATAAGGATGAGATCGCTCTCGTGAACAGCTTCAACGGATTCACTACTGCCTTCACCGGATTCTTCAGAAACAGGGAAAACATGTTCAGCGAAGAGGCCAAGAGCACCTCTATCGCTTTCAGATGCATCAACGAGAACCTCACGCGTTACATCAGCAACATGGACATCTTCGAGAAGGTAACATTCCTTAGTTACCTTTCTTTTCTTTTTCCATCATAAGTTTATAGATTGTACATGCTTTGAGATTTTTCTTTGCAAACAATCTCAGGTGGACGCCATCTTCGATAAGCACGAGGTGCAAGAAATCAAAGAGAAGATCCTCAACAGCGACTACGACGTCGAGGACTTTTTTGAAGGGGAGTTCTTCAACTTCGTTCTCACCCAAGAGGGCATCGACGTGTACAACGCTATTATCGGAGGATTCGTGACCGAGTCTGGGGAGAAGATTAAGGGACTCAACGAGTACATCAACCTGTACAACCAGAAAACGAAGCAGAAGCTCCCGAAGTTCAAGCCGCTCTACAAGCAGGTCTGTCTTTCCTATTTCATATGTTTAATCCTAGGAATTTGATCAATTGATTGTATGTATGTCGATCCCAAGACTTTCTTGTTCACTTATATCTTAACTCTCTCTTTGCTGTTTCTTGCAGGTTCTCTCTGATCGTGAGAGCCTCTCATTTTACGGTGAGGGTTACACCTCTGACGAGGAAGTGCTTGAGGTTTTCCGTAACACCCTCAACAAGAACAGCGAGATCTTCTCGTCCATCAAGAAGTTGGAGAAGCTTTTCAAGAACTTCGACGAGTACAGCAGCGCTGGGATCTTCGTTAAGAACGGACCTGCTATCAGCACCATCAGCAAGGATATTTTCGGCGAGTGGAACGTGATCAGGGACAAGTGGAATGCTGAGTACGATGACATCCACCTCAAGAAGAAGGCTGTCGTCACTGAGAAGTACGAGGATGACAGGCGTAAGTCGTTCAAGAAGATCGGCTCTTTCAGCCTCGAGCAGCTTCAAGAATACGCTGATGCTGATCTCAGCGTGGTCGAGAAGCTCAAAGAGATCATCATCCAGAAGGTCGACGAGATCTACAAGGTAAGTTGTTACTTATGATTGTTTTCCTCTCTGCTACATGTATTTTGTTGTTCATTTCTGTAAGATATAAGAATTGAGTTTTCCTCTGATGATATTATTAGGTGTACGGGTCCTCTGAGAAGTTGTTCGATGCTGATTTCGTCCTCGAGAAGAGTCTGAAGAAGAACGACGCTGTCGTCGCGATCATGAAGGATTTGCTCGACAGCGTGAAGTCCTTCGAGAACTATATCAAGGCCTTCTTCGGAGAGGGCAAAGAGACTAATAGGGACGAGTCTTTCTACGGGGATTTCGTGCTCGCTTACGATATCCTCCTCAAGGTGGACCATATCTACGACGCCATCAGAAACTACGTGACCCAGAAGCCTTACAGCAAGGACAAGTTCAAGTTGTACTTTCAGAACCCGCAGTTCATGGGCGGATGGGACAAAGACAAAGAGACAGATTACAGGGCCACCATCCTCAGGTTAGTATCATATGAAGAAATACCTAGTTTCAGTTGATGAATGCTATTTTCTGACCTCAGTTGTTCTCTTTTGAGAATTATTTCTTTTCTAATTTGCCTGATTTTTCTATTAATTCATTAGGTACGGGTCTAAGTACTACCTGGCCATCATGGACAAGAAATACGCCAAGTGCCTCCAAAAGATCGACAAGGATGACGTGAACGGGAACTATGAGAAGATCAACTACAAGCTCCTTCCGGGACCGAACAAGATGCTTCCTAAGGTGTTCTTCAGCAAGAAATGGATGGCCTACTACAACCCGTCTGAGGACATCCAGAAAATCTACAAGAACGGGACCTTCAAGAAAGGCGACATGTTCAACCTCAACGACTGCCACAAGCTCATCGATTTCTTCAAGGACAGCATCTCGCGTTACCCGAAGTGGTCTAACGCTTACGACTTTAACTTCAGCGAGACAGAAAAGTACAAGGATATCGCCGGGTTCTACCGTGAGGTTGAGGAACAGGGTTACAAAGTTAGCTTCGAGAGCGCCTCCAAGAAAGAGGTAAATCCTGGTCCACACTTTTACGATAAAAACACAAGATTTTAAACTATGAACTGATCAATAATCATTCCTAAAAGACCACACTTTTGTTTTGTTTCTAAAGTAATTTTTACTGTTATAACAGGTTGACAAGTTGGTCGAAGAGGGCAAGCTCTACATGTTCCAGATCTATAACAAGGACTTCTCCGACAAGAGCCACGGAACTCCTAACCTCCATACGATGTACTTCAAGCTGCTTTTCGACGAGAACAACCACGGGCAGATCAGACTTTCTGGTGGTGCTGAACTCTTCATGCGTAGGGCCTCACTCAAGAAAGAAGAGTTGGTTGTTCACCCGGCCAACTCTCCAATCGCTAACAAGAATCCTGACAACCCGAAAAAGACCACCACGCTGTCTTACGACGTCTACAAGGACAAAAGGTTCAGCGAGGACCAGTACGAGCTTCATATCCCGATCGCTATCAACAAGTGCCCGAAGAACATCTTCAAGATCAATACCGAGGTAAGGACTTCTCATGAATATTAGTGGCAGATTAGTGTTGTTAAAGTCTTTGGTTAGATAATCGATGCCTCCTAATTGTCCATGTTTTACTGGTTTTCTACAATTAAAGGTGAGGGTGCTGCTCAAGCACGATGATAACCCTTACGTGATCGGAATCGATCGTGGTGAGAGAAACCTCCTCTACATCGTTGTGGTGGACGGAAAGGGAAACATCGTCGAGCAGTACAGCCTGAACGAGATTATCAACAATTTCAACGGCATCAGGATCAAGACCGACTACCACTCACTCCTCGATAAGAAAGAAAAAGAGCGTTTCGAGGCCAGGCAGAACTGGACTTCTATCGAAAACATCAAAGAGTTGAAGGCCGGCTACATCTCTCAGGTGGTGCATAAGATCTGCGAGCTGGTGGAAAAGTACGATGCTGTGATCGCTCTTGAGGACCTCAACTCTGGGTTCAAGAACAGTAGAGTGAAGGTAAGTTCTGCATTTGGTTATGCTCCTTGCATTTTAGGTGTTCGTCGCACTTCCATTTCCATGAATAGCTAAGATTTTTTTTCTCTGCATTCATTCTTCTTGCCTCAGTTCTAACTGTTTGTGGTATTTTTGTTTTAATTATTGCTACAGGTTGAGAAGCAGGTCTACCAAAAGTTCGAGAAGATGCTCATCGACAAGCTCAACTACATGGTGGACAAAAAGAGCAACCCTTGCGCTACCGGTGGTGCTCTTAAGGGATACCAGATCACGAACAAGTTCGAGTCCTTCAAGAGCATGAGCACCCAGAACGGCTTCATCTTCTATATCCCTGCTTGGCTCACCAGCAAGATCGATCCTTCTACTGGTTTCGTGAACCTGCTCAAGACCAAGTACACCTCGATCGCCGACAGCAAGAAGTTCATCTCGTCTTTCGACAGGATCATGTACGTGCCGGAAGAGGATCTTTTCGAGTTCGCTCTCGACTATAAGAACTTCAGCAGGACCGACGCCGACTACATTAAGAAGTGGAAGCTCTACTCCTACGGGAACCGTATCAGGATCTTCCGAAATCCGAAGAAAAACAACGTGTTCGACTGGGAAGAAGTGTGCCTCACCTCTGCCTACAAAGAACTGTTCAACAAGTACGGCATCAACTACCAGCAGGGTGATATCAGGGCTCTTTTGTGCGAGCAGAGCGACAAGGCATTCTACAGCTCATTCATGGCCCTCATGTCTCTCATGCTCCAGATGAGGAACTCTATCACCGGAAGGACCGATGTGGACTTCCTTATCTCTCCGGTCAAGAACTCTGACGGGATCTTCTACGACAGCCGTAACTATGAGGCTCAAGAGAACGCTATCCTGCCGAAGAATGCTGATGCAAACGGGGCTTACAACATTGCGAGAAAGGTAAAGCAACTGTGTTTTAATCAATTTCTTGTCAGGATATATGGATTATAACTTAATTTTTGAGAAATCTGTAGTATTTGGCGTGAAATGAGTTTGCTTTTTGGTTTCTCCCGTGTTATAGGTTCTCTGGGCTATCGGGCAGTTTAAGAAAGCGGAAGATGAGAAGCTGGACAAGGTGAAGATCGCCATCTCCAACAAAGAGTGGCTTGAGTACGCTCAGACCTCCGTTAAGCACAAGAGGCCTGCTGCTACTAAGAAAGCTGGTCAGGCTAAGAAGAAGAAATGA

V1 array

GACCAAGCCCGTTATTCTGACAGTTCTGGTGCTCAACACATTTATATTTATCAAGGAGCACATTGTTACTCACTGCTAGGAGGGAATCGAACTAGGAATATTGATCAGAGGAACTACGAGAGAGCTGAAGATAACTGCCCTCTAGCTCTCACTGATCTGGGTCGCATAGTGAGATGCAGCCCACGTGAGTTCAGCAACGGTCTAGCGCTGGGCTTTTAGGCCCGCATGATCGGGCTTTTGTCGGGTGGTCGACGTGTTCACGATTGGGGAGAGCAACGCAGCAGTTCCTCTTAGTTTAGTCCCACCTCGCCTGTCCAGCAGAGTTCTGACCGGTTTATAAACTCGCTTGCTGCATCAGACTTGTAATTTCTACTAAGTGTAGATNNNNNNNNNNNNNNNNNNNNNNNTAATTTCTACTAAGTGTAGATNNNNNNNNNNNNNNNNNNNNNNNTAATTTCTACTAAGTGTAGATNNNNNNNNNNNNNNNNNNNNNNNTAATTTCTACTAAGTGTAGATNNNNNNNNNNNNNNNNNNNNNNNTAATTTCTACTAAGTGTAGATGTCCCTTCGAAGGGCAATTCTGCAGATATCCATCACACTGGCGGCCGCTCGAGGTCGAGGGTATCGATAAGCTTTTTTTTTTT

V2 array

GACCAAGCCCGTTATTCTGACAGTTCTGGTGCTCAACACATTTATATTTATCAAGGAGCACATTGTTACTCACTGCTAGGAGGGAATCGAACTAGGAATATTGATCAGAGGAACTACGAGAGAGCTGAAGATAACTGCCCTCTAGCTCTCACTGATCTGGGTCGCATAGTGAGATGCAGCCCACGTGAGTTCAGCAACGGTCTAGCGCTGGGCTTTTAGGCCCGCATGATCGGGCTTTTGTCGGGTGGTCGACGTGTTCACGATTGGGGAGAGCAACGCAGCAGTTCCTCTTAGTTTAGTCCCACCTCGCCTGTCCAGCAGAGTTCTGACCGGTTTATAAACTCGCTTGCTGCATCAGACTTGAAATTACTGATGAGTCCGTGAGGACGAAACGAGTAAGCTCGTCTAATTTCTACTAAGTGTAGATNNNNNNNNNNNNNNNNNNNNNNNGGCCGGCATGGTCCCAGCCTCCTCGCTGGCGCCGGCTGGGCAACATGCTTCGGCATGGCGAATGGGACTTTTTGACCAAGCCCGTTATTCTGACAGTTCTGGTGCTCAACACATTTATATTTATCAAGGAGCACATTGTTACTCACTGCTAGGAGGGAATCGAACTAGGAATATTGATCAGAGGAACTACGAGAGAGCTGAAGATAACTGCCCTCTAGCTCTCACTGATCTGGGTCGCATAGTGAGATGCAGCCCACGTGAGTTCAGCAACGGTCTAGCGCTGGGCTTTTAGGCCCGCATGATCGGGCTTTTGTCGGGTGGTCGACGTGTTCACGATTGGGGAGAGCAACGCAGCAGTTCCTCTTAGTTTAGTCCCACCTCGCCTGTCCAGCAGAGTTCTGACCGGTTTATAAACTCGCTTGCTGCATCAGACTTGAAATTACTGATGAGTCCGTGAGGACGAAACGAGTAAGCTCGTCTAATTTCTACTAAGTGTAGATNNNNNNNNNNNNNNNNNNNNNNNGGCCGGCATGGTCCCAGCCTCCTCGCTGGCGCCGGCTGGGCAACATGCTTCGGCATGGCGAATGGGACTTTTTTTACGGCGGCAGGGAGAGTTTTAACATTGACTAGCGTGCTGATAATTTGTGAGAAATAATAATTGACAAGTAGATACTGACATTTGAGAAGAGCTTCTGAACTGTTATTAGTAACAAAAATGGAAAGCTGATGCACGGAAAAAGGAAAGAAAAAGCCATACTTTTTTTTAGGTAGGAAAAGAAAAAGCCATACGAGACTGATGTCTCTCAGATGGGCCGGGATCTGTCTATCTAGCAGGCAGCAGCCCTACCAACCTCACGGGCCAGCAATTACGAGTCCTTCTAAAACGTCCCGCCGAGGGCGCGTGGCCGTGCTGTGCAGCAGCACGTCTAACATTAGTCCCACCTCGCCAGTTTACAGGGAGCAGAACCAGCTTATAAGCGGAGGCGCGGCACCAAGAAGCAAAATTACTGATGAGTCCGTGAGGACGAAACGAGTAAGCTCGTCTAATTTCTACTAAGTGTAGATNNNNNNNNNNNNNNNNNNNNNNNGGCCGGCATGGTCCCAGCCTCCTCGCTGGCGCCGGCTGGGCAACATGCTTCGGCATGGCGAATGGGACTTTTTGGCGGCAGGGAGAGTTTTAACATTGACTAGCGTGCTGATAATTTGTGAGAAATAATAATTGACAAGTAGATACTGACATTTGAGAAGAGCTTCTGAACTGTTATTAGTAACAAAAATGGAAAGCTGATGCACGGAAAAAGGAAAGAAAAAGCCATACTTTTTTTTAGGTAGGAAAAGAAAAAGCCATACGAGACTGATGTCTCTCAGATGGGCCGGGATCTGTCTATCTAGCAGGCAGCAGCCCTACCAACCTCACGGGCCAGCAATTACGAGTCCTTCTAAAACGTCCCGCCGAGGGCGCGTGGCCGTGCTGTGCAGCAGCACGTCTAACATTAGTCCCACCTCGCCAGTTTACAGGGAGCAGAACCAGCTTATAAGCGGAGGCGCGGCACCAAGAAGCAAAATTACTGATGAGTCCGTGAGGACGAAACGAGTAAGCTCGTCTAATTTCTACTAAGTGTAGATNNNNNNNNNNNNNNNNNNNNNNNGGCCGGCATGGTCCCAGCCTCCTCGCTGGCGCCGGCTGGGCAACATGCTTCGGCATGGCGAATGGGACTTTTT

Guide sequences given as arrayed order according to figure 1:

HORVU.MOREX.r3.2HG0184740

1. TGGAGGATGGCGGCGTCGTGGGC
2. TCAGTAAGCATGTTCAAGAACGG
3. AGCGGGGCAGACGTGCTGGGGCG
4. ATGCTAAAAAACACTCGACCACG

HORVU.MOREX.r3.6HG0611290

1. TCACACGAACTGCCGGAACGTTG
2. AAGGACATGATCATGGAGGGTGA
3. GACCTACAAGACCTACAAGTGTG
4. ATTCATATAAGCTTGTACTACTT

HORVU.MOREX.r3.7HG0640970

1. TGAACCCCATCATCTCCATGCTC
2. TGTTCTTCCTTGACCAGCATCAT
3. GGAACTATTTCAGTCAATCATCG
4. CTCGAAAGGAGGAAGTGTCAAGA

HORVU.MOREX.r3.2HG0133680

1. CCATAGCTCCTCCTTGAGTCGCT
2. AGGCTGAGAGATGGTGCGCCATT
3. TTCTGACCTGATGTGGAGTTCTG
4. GCAGTGAAGAGGAAAATTAAACG

HORVU.MOREX.r3.1HG0069960

1. TCCATAGTGAGAAGAGGTGTGAG
2. TCGCGACCTGGGTCTTTCCTTCA
3. GTGCTGCACAATGTCAACAACTG
4. ACCCCTGCTGCATCCCCATCATC
